# MSX2 safeguards syncytiotrophoblast fate of human trophoblast stem cells

**DOI:** 10.1101/2021.02.03.429538

**Authors:** Ruth Hornbachner, Andreas Lackner, Sandra Haider, Martin Knöfler, Karl Mechtler, Paulina A. Latos

## Abstract

The majority of placental pathologies are associated with failures in trophoblast differentiation, yet the underlying transcriptional regulation is poorly understood. Here, we use human trophoblast stem cells to elucidate the function of the transcription factor MSX2 in trophoblast specification. We show that depletion of MSX2 de-represses the syncytiotrophoblast program, while forced expression of MSX2 blocks it. We demonstrate that a large proportion of the affected genes are directly bound and regulated by MSX2 and identify components of the SWI/SNF complex as its strong interactors. Our findings uncover the pivotal role of MSX2 in cell fate decisions that govern human placental development and function.

## Introduction

The placenta is a vital organ that sustains mammalian development *in utero.* It provides an interface for gas, nutrients and metabolite exchange and serves hormonal as well as immunological functions. The human placenta comprises of three major trophoblast cell types: i) the extravillous trophoblast (EVT) that invades maternal decidua and remodels spiral arteries, ii) the multinuclear syncytiotrophoblast (ST) that provides the site of exchange and produces placental hormones and iii) the cytotrophoblast (CT), a multipotent progenitor population that gives rise to both EVT and ST (Supplemental Fig.S1A) (Turco and Moffett 2019). Precise and coordinated differentiation of these cell types is a prerequisite for a successful pregnancy. Failures in cell fate commitment as well as impaired trophoblast development and function may lead to placental disorders including foetal growth restriction, miscarriage and preeclampsia (Brosens et al. 2011). However, the underlying molecular causes of these pathologies remain largely unknown.

Placental growth and development are orchestrated by the spatially and temporarily coordinated actions of various transcription factors (TF). Despite the fact that murine and human placentas exert equivalent functions, they are morphologically different, and their governing TF networks only partially overlap. While TEAD4, GATA3, CDX2, ELF5 and TFAP2C are expressed in both mouse and human TSCs and their respective *in vivo* counterparts, Eomes and MSX2 are expressed specifically in mouse and human, respectively (Hemberger et al. 2020; Latos and Hemberger 2016; Liang et al. 2016). Both mouse and human placentas comprise syncytiotrophoblast, that results from fusion of progenitor cells and constitutes the direct mother/embryo interface. However, the molecular mechanisms, in particular the TF networks, driving its development and function are species-specific and remain elusive (Turco and Moffett 2019; Latos and Hemberger 2016).

Recent establishment of human trophoblast stem cells (hTSCs) from CT have the potential to revolutionize trophoblast research. Human TSC self-renew and are multipotent, as they have the ability to differentiate into both ST and EVT. Overall, global gene expression, DNA methylation profiling and functional analysis have demonstrated that hTSCs and their *in vitro* ST and EVT derivatives faithfully recapitulate the *in vivo* counterparts and provide an excellent model to study molecular mechanisms driving human placental development and disease (Okae et al. 2018).

Here, we addressed the placental function of MSX2 (msh homeobox 2) TF using the hTSC *in vitro* model. We demonstrated that levels of MSX2 determine trophoblast cell identity as its depletion results in precocious ST differentiation, while its forced expression blocks it. We found that a significant proportion of genes affected upon these genetic manipulations are bound by MSX2, suggesting direct regulation. To further dissect how MSX2 exerts its function, we identified its interactome and revealed components of the SWI/SNF complex as major interactors of MSX2. Taken together, our findings established MSX2 as a central regulator of cell fate decisions in human trophoblast and provide novel molecular insights into its mode of action.

## Results and Discussion

### Depletion of MSX2 results in loss of silencing of syncytiotrophoblast genes

Our previous RNA profiling revealed that the TF MSX2 is highly expressed in CT and down-regulated upon ST differentiation *in vitro,* prompting us to address its role in human trophoblast (Haider et al. 2018). First, we examined expression of MSX2 in the 1^st^ trimester human placenta and showed that it was readily expressed in CT but not in the ST compartment, as indicated by co-staining with the ST marker chorionic gonadotropin beta (CGB) (Fig.1A). Similarly, MSX2 was highly expressed in hTSCs and rapidly downregulated upon ST differentiation induced by 2μM forskolin (Supplemental Fig.S1B), demonstrating hTSCs as a reliable model to address the role of MSX2 in human placenta. To determine its function, we attempted to knock-out MSX2 in hTSCs, but were unable to retrieve clones, possibly due to the essential role of MSX2. Therefore, we depleted MSX2 using two different shRNAs (MSX2_KD-1 and MSX2_KD-2) along a control shRNA (CTRL), by lentiviral transduction. MSX2 transcript levels were reduced in the MSX2_KD-1 and MSX2_KD-2 lines by up to 80% and these results were confirmed on the protein level (Fig.1B,C). Strikingly, while the control cells formed tight, epithelial colonies, the MSX2_KDs gradually lost the hTSC morphology and proliferative capacity, indicating spontaneous differentiation (Fig.1D). Expression analysis revealed a substantial upregulation of ST markers including *CGB, GCM1* and *ENDOU* in culture conditions supporting hTSC self-renewal (SR) (Fig.1B,C). Importantly, this effect was specific to MSX2 depletion, as doxycycline-inducible expression of mouse Msx2 coding sequence, which is resistant to the shRNAs targeting human MSX2, fully rescued the MSX2_KD phenotype (Supplemental Fig.S1C,D). To gain a global overview of gene expression changes caused by depletion of MSX2, we performed RNA-seq analysis on MSX2_KD-1, MSX2_KD-2 and CTRL cells. We identified 152 down-regulated and 522 up-regulated genes (cut off: ļlog2FCļ>1, p adj<0.05), in line with the reported function of MSX2 as a transcriptional repressor (Catron et al. 1995) (Fig.1E and Supplemental Fig.S1 E). Among the up-regulated genes we noted numerous ST markers including *CGA*, *CGB1(2,3,5,7,8), LHB, PSG1(3,4,6,8,9), CSH1, CSH2, ERVV-1, ERVV-2, INSL4, SDC1, HSD11B2, GPR78, LGALS14, SLC6A2,* and *CYP19A1* (Fig.1E and Supplemental Fig.S1F). Importantly, their expression levels were comparable to the *in vitro* differentiated ST (Supplemental Fig.S1F). Gene set enrichment analysis (GSEA) revealed enrichment in terms relating to hormone activity, endocrine and hormone metabolic processes, and cofactor transport (Supplemental Fig.S1G). Taken together, MSX2 depletion results in the massive loss of silencing of ST genes, suggesting that MSX2 prevents access to ST fate. Previous reports demonstrated that Msx2/MSX2 deficiency results in impaired osteogenesis, chondrogenesis and in defective tooth, hair follicle and mammary gland development (Satokata et al. 2000; Babajko et al. 2014; Wilkie et al. 2000). A common theme of these diverse phenotypes was defective progenitor proliferation and their imbalanced differentiation, consistent with our observations that depletion of MSX2 in hTSCs resulted in precocious ST differentiation. These findings have important implications as the underlying causes of various placental pathologies, including severe forms of foetal growth restriction and preeclampsia have their roots in defective ST development and function.

**Figure 1.**
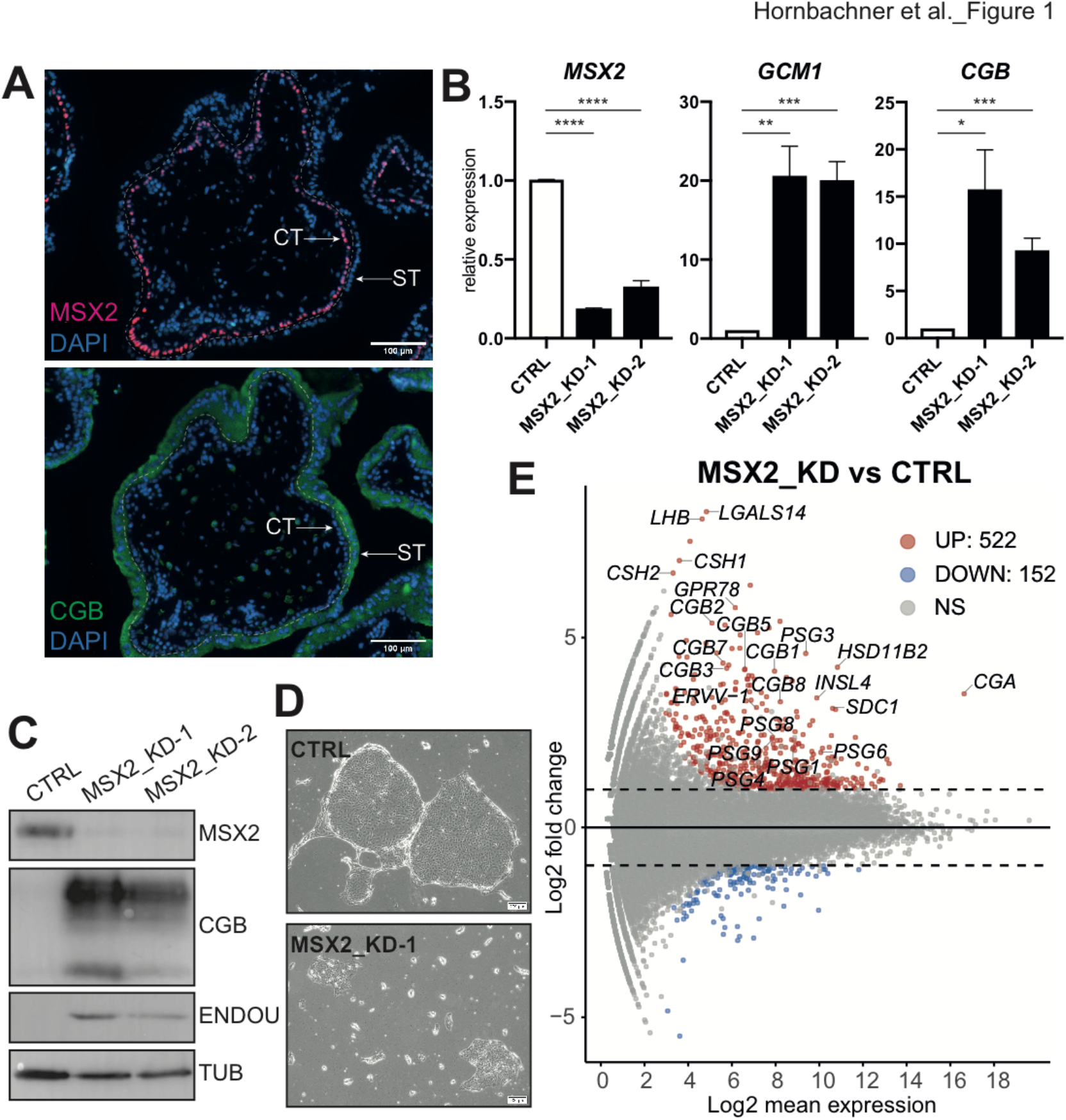
Depletion of MSX2 results in loss of silencing of syncytiotrophoblast genes (*A*) Immunofluorescence staining of a first trimester human placenta for MSX2 and CGB, counterstained with DAPI. ST: syncytiotrophoblast, CT: cytotrophoblast, dashed line separates ST from CT. *(B-C)* RTQPCR *(B)* Western blot (*C*) expression and analysis of control (CTRL) and MSX2-depleted (MSX2_KD1 and MSX2_KD2) hTSC lines cultured in stem cell conditions for MSX2 and syncytiotrophoblast markers GCM1, CGB and ENDOU. (*B*) The bars represent a mean of four (n=4) biological replicates with S.E.M. ****p<0.0001, ***p<0.001, **p<0.01, *p<0.05, ns: not significant. (*C*) Tubulin (TUB) serves as a loading control. *(D)* Phase contrast microscope images of hTSC 10 days after viral transduction with MSX2_KD1 or control (CTRL) constructs. *(E)* MA plot from differentially expressed genes between MSX2-depleted (MSX2_KD1 (n=3) and MSX2_KD2 (n=1)) and control (CTRL (n=3)). Analysis (cut-off: |log2FC|>1, p adj <0.05) revealed 522 up-regulated and 152 down-regulated genes in MSX2_KD lines compared to control. Significantly up- (UP) and downregulated (DOWN) genes are indicated in red and blue, respectively. Dashed black line highlights |log2FC|>1. NS: not significant.

### Ectopic expression of MSX2 blocks syncytiotrophoblast cell fate

We showed that depletion of MSX2 leads to de-repression of ST genes, supporting the role of MSX2 as a transcriptional repressor. Based on this result, we reasoned that forced expression of MSX2 would block ST differentiation. To test this hypothesis, we cloned a flag-tagged human MSX2 protein coding sequence under the control of a doxycycline-inducible promoter, generated stable hTSC lines (MSX2_iOX) (Supplemental Fig.S2A) and concomitantly induced ST differentiation and ectopic expression of MSX2 using forskolin and doxycycline, respectively. After 72h of treatment we observed a robust formation of syncytia and CGB expression in non-treated cells and lack thereof in dox-treated, MSX2-expressing cells (Fig.2A). Instead, MSX2-expressing cells formed large, flattened colonies with clearly visible cell borders (Supplemental Fig.S2C). Gene expression analysis revealed induction of ST markers including *GCM1, SDC1, ERVW-1, ENDOU* and *CGB* in control, but not in the dox-treated MSX2_iOX cells (Fig.2B and Supplemental Fig.S2C). To examine the global gene expression changes, we performed RNA-seq on untreated and dox-treated MSX2_iOX cells cultured in SR and for 6 days in ST conditions. Our analysis confirmed down-regulation of selfrenewal markers *(TEAD4, TP63, ELF5)* upon ST induction in both -dox and +dox. Interestingly, a massive up-regulation of a variety of ST markers including *CGA*, *CGB1(2,3,5,7,8), PSG1(3,4,6,8,9), ERVV-1*, *ERVV-2, GCM1, SDC1, CYP26A1, HSD11B2* and others was only observed in -dox but not in +dox treatment (Fig.2C and Supplemental Fig.S2D,E). We identified 1285 up-regulated and 1433 down-regulated genes in +dox vs -dox MSX2_iOX ST cells (cut off: |log2FC|>1, p adj<0.05), indicating that MSX2 acts as a strong transcriptional repressor preventing up-regulation of ST genes (Fig.2C and Supplemental Fig.S2D). Indeed, the GSEA analysis of the de-regulated genes confirmed enrichment in terms related to cell cycle and depletion of terms related to hormone metabolism (Supplemental Fig.S2F). Since we demonstrated that depletion of MSX2 led to spontaneous ST differentiation in SR conditions and conversely, forced expression of MSX2 prevented ST differentiation, we sought to identify a shared set of mis-regulated genes. Based on the RNA-seq datasets, we compared genes that were up-regulated upon MSX2 depletion in SR conditions to those that were down-regulated upon MSX2 forced expression in ST conditions and revealed that 239 genes were shared, i.e. commonly mis-regulated (Fig.2D). These results demonstrate that MSX2 acts as a strong transcriptional repressor that presides over a network of ST-specific genes and regulates key cell fate decisions during trophoblast differentiation.

**Figure 2.**
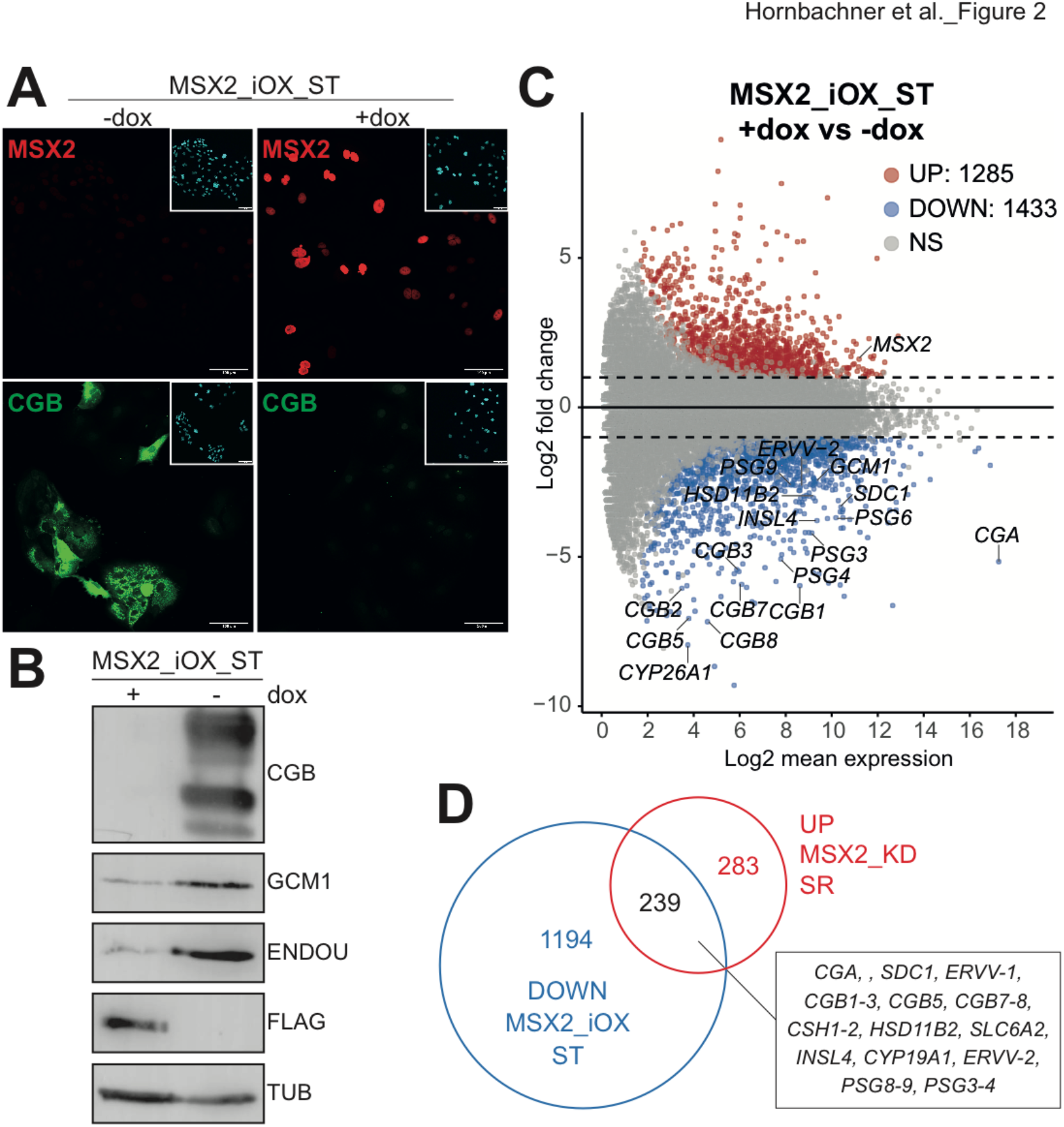
Ectopic expression of MSX2 blocks syncytiotrophoblast cell fate (*A*) Immunofluorescence staining for MSX2 and CGB in hTSC line carrying a doxycycline (dox) inducible MSX2 transgene (MSX2_iOX) and differentiated to syncytiotrophoblast (ST) in the presence (+dox) or absence (-dox) of doxycycline. (*B*) Western blot analysis for syncytiotrophoblast markers CGB, GCM1 and ENDOU in hTSC line carrying a doxycycline (dox) inducible MSX2 transgene and differentiated to syncytiotrophoblast (ST) in the presence (+dox) or absence (-dox) of dox. MSX2 transgene carries a 3xFlag-tag; tubulin was used as loading control. (*C*) MA plot from differential expressed genes between MSX2_iOX ST +dox (n=2) and MSX2_iOX ST -dox (n=3). Analysis (cut-off: |log2FC|>1, p adj <0.05) revealed 1285 up-regulated and 1433 down-regulated genes in MSX2_iOX ST +dox compared to MSX2_iOX ST-dox. Significantly up- (UP) and downregulated (DOWN) genes are indicated in red and blue, respectively. Dashed black line highlights log fold changes of −2 and 2. NS: not significant. (*D*) Venn diagram showing overlap of genes that were up-regulated upon MSX2 depletion (MSX2_KD) in self-renewal (SR) conditions (compared to KD control) and genes that were down-regulated (i.e. did not get upregulated) upon induced expression of MSX2 transgene (MSX2_iOX) during syncytiotrophoblast differentiation (ST) (compared to -dox control). The Venn diagram is based on the RNA-seq analysis detailed in Fig.1E and 2C.

Our findings that forced expression of MSX2 blocks ST differentiation are in line with previous reports showing that forced expression of mouse Msx1 and Msx2 negatively regulates differentiation in multiple mesenchymal and epithelial cell types including muscle, adipocytes, cartilage, bone and mammary gland epithelium (Hu et al. 2001). Consistent with its function as an inhibitor of differentiation, increased expression of MSX2 was observed in a number of cancers and correlated with tumour invasiveness (Satoh et al. 2012). The role of MSX2 as a strong determinant of cell identity was also demonstrated in human embryonic stem cells (hESCs), where forced expression of MSX2 resulted in precocious mesendoderm differentiation and conversely, MSX2 deficient hESCs were severely impaired in mesendodermal differentiation (Wu et al. 2015). Likewise, MSX2 was reported to initiate and accelerate a molecular program driving mesenchymal stem/stromal cell specification (Zhang et al. 2018). Taken together, these observations suggest, that independent of the context, expression of MSX2/Msx2 marks a transient population of multipotent, proliferative progenitors, that, as development progresses and MSX2/Msx2 expression vanishes, will further differentiate into specialized cell types.

### MSX2 binds and silences syncytiotrophoblast genes in hTSCs

To explore which of the genes that are mis-regulated upon genetic manipulations of MSX2, are directly bound and regulated by MSX2, we performed chromatin immunoprecipitation followed by high-throughput sequencing (ChIP-seq) in SR hTSC. We uncovered 2261 MSX2 binding sites, of which nearly 30% were located in promoters, first exons, 5’UTRs and first introns of genes (Fig.3A). Next, we overlaid genes that were up- and down-regulated upon MSX2 depletion with 1908 annotated genes bound by MSX2 and revealed that over 17.4% (*p*-value 7.842E-14) of the up-regulated genes in contrast to 9.2% (*p*-value 0.271) of the down-regulated genes are direct targets of MSX2 (Fig.3B and Supplemental Fig.S3A). This result further supports the notion that MSX2 acts predominantly as a transcriptional repressor. Additionally, we overlapped MSX2 targets with genes that were up- and down-regulated upon forced MSX2 expression during ST differentiations, and showed that 13% (*p*-value 1.050E-14) and 15% (*p*-value 6.753E-28) respectively, are bound by MSX2 in hTSCs (Fig.3C and Supplemental Fig.S3B). To further narrow down the direct, functional targets of MSX2, we overlapped genes that were bound by MSX2 and either up-regulated upon MSX2 depletion or down-regulated upon forced MSX2 expression during ST differentiation. The shared subset of 43 genes (*p*-value 7.86E-68) contained predominantly genes associated with ST including the *PSG* family, *INSL4, SDC1, TBX3, SLC6A4, SLC6A2, SLC40A1, AKR1B1, NUCB2, LGALS13, PRKCZ* and *IL1R1* (Fig.3D,E). These genes are direct targets of MSX2 and their functional relevance for ST differentiation has yet to be determined.

**Figure 3.**
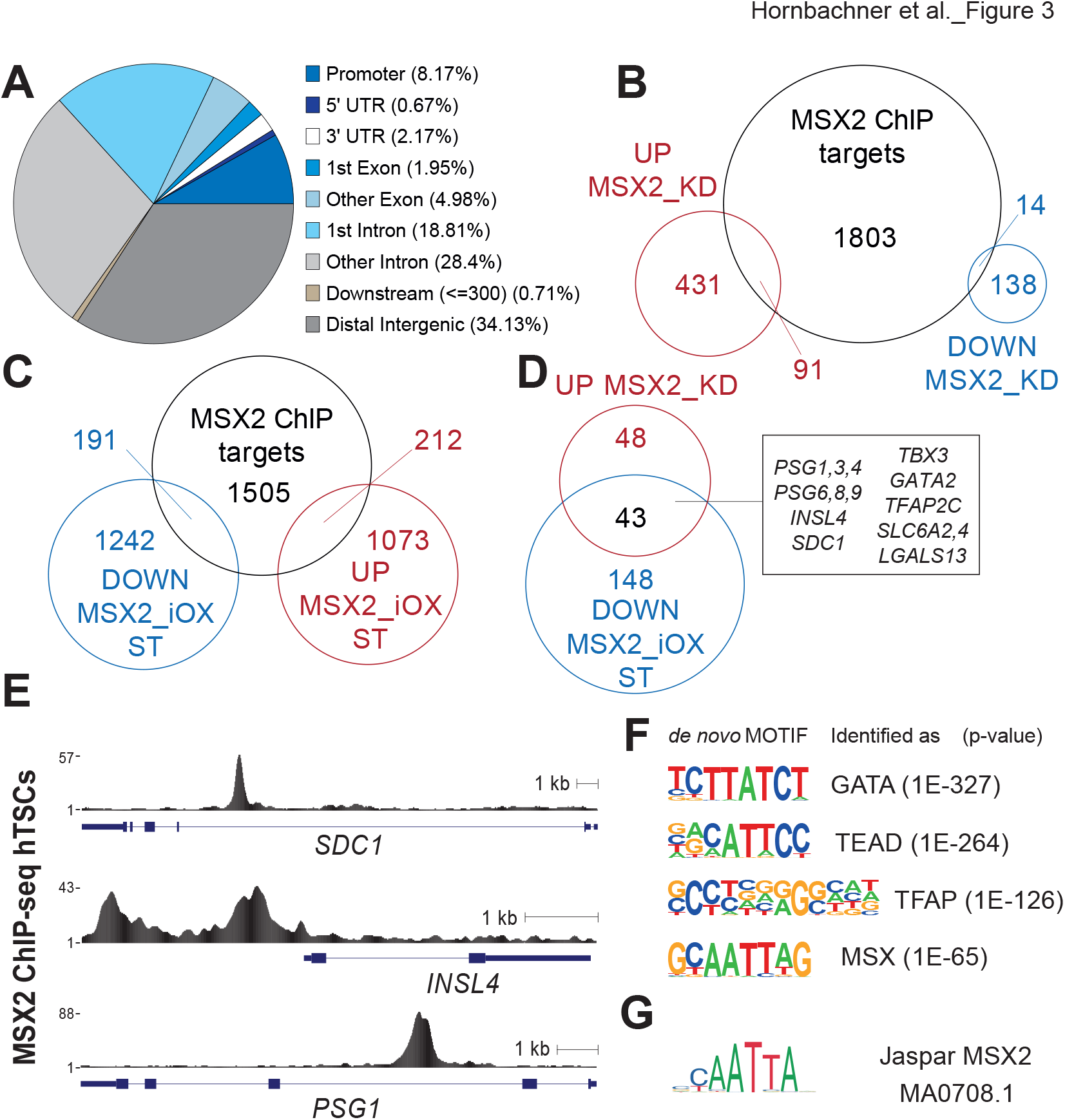
MSX2 binds and silences syncytiotrophoblast genes in hTSC (*A*) Proportion of MSX2 ChIP-seq peaks overlapping genomic features in hTSC. (*B*) Venn diagram depicting overlap between genes bound by MSX2 (identified by ChIP-seq) and genes that were either down-regulated (MSX2KD_down) or up-regulated (MSx2KD_down) upon MSX2 depletion (based on RNA-seq analysis). (*C*) Venn diagram depicting overlap between genes bound by MSX2 (identified by ChIP-seq) and genes that were either down-regulated (MSX2_iOXST_down) or up-regulated (MSX2_iOXST_down) upon forced MSX2 expression during syncytiotrophoblast (ST) differentiation (based on RNA-seq analysis). (*D*) Venn diagram showing overlap between genes that are bound by MSX2 and downregulated upon forced MSX2 expression during syncytiotrophoblast (ST) differentiation (MSX2_iOXST_down) and those bound by MSX2 and up-regulated upon MSX2 depletion. (*E*) RPKM normalized MSX2 binding profiles at selected loci identified in D. (*F*) *De novo* motifs identified by HOMER. (*G*) Matching of *de novo* motif similar to MSX2 with the deposited MSX2 motif from Jaspar.

To gain better molecular insights into the MSX2-driven gene regulation, we performed *de novo* motif analysis using HOMER. The top 4 identified motifs shared high similarity with the GATA, TEAD, TFAP, and MSX motifs (Fig.3F). Comparison of the *de novo* motif for MSX with published MSX2 motifs (Ma et al. 1996) confirmed its identity (Fig.3G). Moreover our data suggest the extension of the currently described human MSX2 motif (Ma et al. 1996) by a flanking G/C nucleotide on each side (Fig.3F,G). A specific search identified MSX2 *de novo* motif instances (Fig.3F) in 69% of the significantly reproducible MSX2 peaks (*p*-value 1E-9, Supplemental Fig.S3C) further validating the motif search and supporting the specificity of the ChIP-seq data. Moreover, known motifs for GATA3, TEAD4 and TFAP2C were strongly enriched in these MSX2 peaks (Supplemental Fig.S3D), suggesting a potential cooperation between those factors and MSX2. Several GATA, TFAP2 and TEAD TF family members are key trophoblast regulators in both mouse and human (Hemberger et al. 2020). TFAP2C, TFAP2A, GATA3 and GATA2 were reported to cooperatively regulate early trophoblast specification *in vitro* (Krendl et al. 2017). Similarly, TEAD4 is a crucial controller of CT identity as its depletion in hTSC resulted in loss of self-renewal and differentiation (Saha et al. 2020). Taken together, our findings highlight MSX2 as a key regulator within the TF network of human trophoblast development.

### MSX2 cooperates with components of the SWI/SNF complex

To bring about transcriptional silencing, transcriptional repressors usually cooperate with large protein complexes including the SWI/SNF, NCoR or NuRD complex as well as other TFs. To gain new molecular insights into the repressive function of MSX2, we set out to determine the MSX2 interactome using rapid immunoprecipitation mass spectrometry of endogenous proteins (RIME) (Mohammed et al. 2016). Using this unbiased protein identification approach, we uncovered MSX2 in addition to numerous high-confidence interaction partners (Fig.4A and Supplemental Fig.S4A). Among them were the key trophoblast regulators GATA3 and TFAP2C. These interactions, together with our previous findings that both GATA3 and TFAP2C DNA binding motifs are overrepresented in the MSX2 ChIP-Seq peaks, raise an exciting possibility that these three factors closely cooperate and co-ordinately regulate transcriptional outputs in hTSC. Strikingly, we also identified numerous components of the mammalian SWI/SNF complex as robust MSX2 interactors: SMARCA4, SMARCA2, SMARCC2, SMARCC1, SMARCB1, SMARCE1, ARID1A, SMARCD1, SMARCD2, DPF2 and ACTL6A, but not other protein complexes (Fig.4A and Supplemental Fig.S4A). We confirmed several of these interactions by co-immunoprecipitation followed by Western blot analysis (Fig.4B). SWI/SNF is a chromatin remodelling complex involved in regulation of gene expression. Upon chromatin recruitment by transcription factors and other proteins, the SWI/SNF complex shifts nucleosomes along the DNA and modifies its accessibility, resulting in a context-dependent transcriptional activation or repression. Importantly, the mammalian SWI/SNF complex exists in multiple forms that are characterized by different subunit compositions. Resulting heterogeneity ensures functional and context-dependent diversity in driving lineage-specific gene expression programs as exemplified by the unique, stagespecific SWI/SNF complexes found in embryonic stem (ES) cells, neural progenitors, postmitotic neurons, cardiac lineage and in cancer (Son and Crabtree 2014; Hota et al. 2019). Based on the identified subunits (SMARCA4, SMARCA2, SMARCC2, SMARCC1, SMARCB1, SMARCE1, ARID1A, SMARCD1, SMARCD2, DPF2 and ACTL6A), we conclude that MSX2 interacts with a canonical version of the BAF complex in hTSC. Importantly, alternative variants (paralogs) exist for several of the subunits and only one of them can be incorporated to the complex at a given polymorphic position. For instance, we detected both SMARCA4 (BRG1) and SMARCA2 (BRM), although they are incorporated into the SWI/SNF complex in a mutually exclusive manner, raising the possibility that several different subcomplexes operate in hTSC. The subunit composition of the MSX2-interacting SWI/SNF complex is similar to the one operating in hESC and distinctive from the one in mESC (Zhang et al. 2014; Ho et al. 2009). Interestingly, in both ESC contexts, the SWI/SNF complex is critical for maintenance of pluripotency and self-renewal, by keeping developmental genes silent and fine-tuning expression of self-renewal genes (Zhang et al. 2014; Ho et al. 2009). Based on our results, we propose that in the context of MSX2, the transcriptional repressor interacts with the SWI/SNF complex, recruits it to a subset of ST-specific genes and confers transcriptional silencing. Overall, while our work provides first insights into the composition and interactions of the SWI/SNF complex in hTSC, extensive studies are required for its full functional characterization.

**Figure 4.**
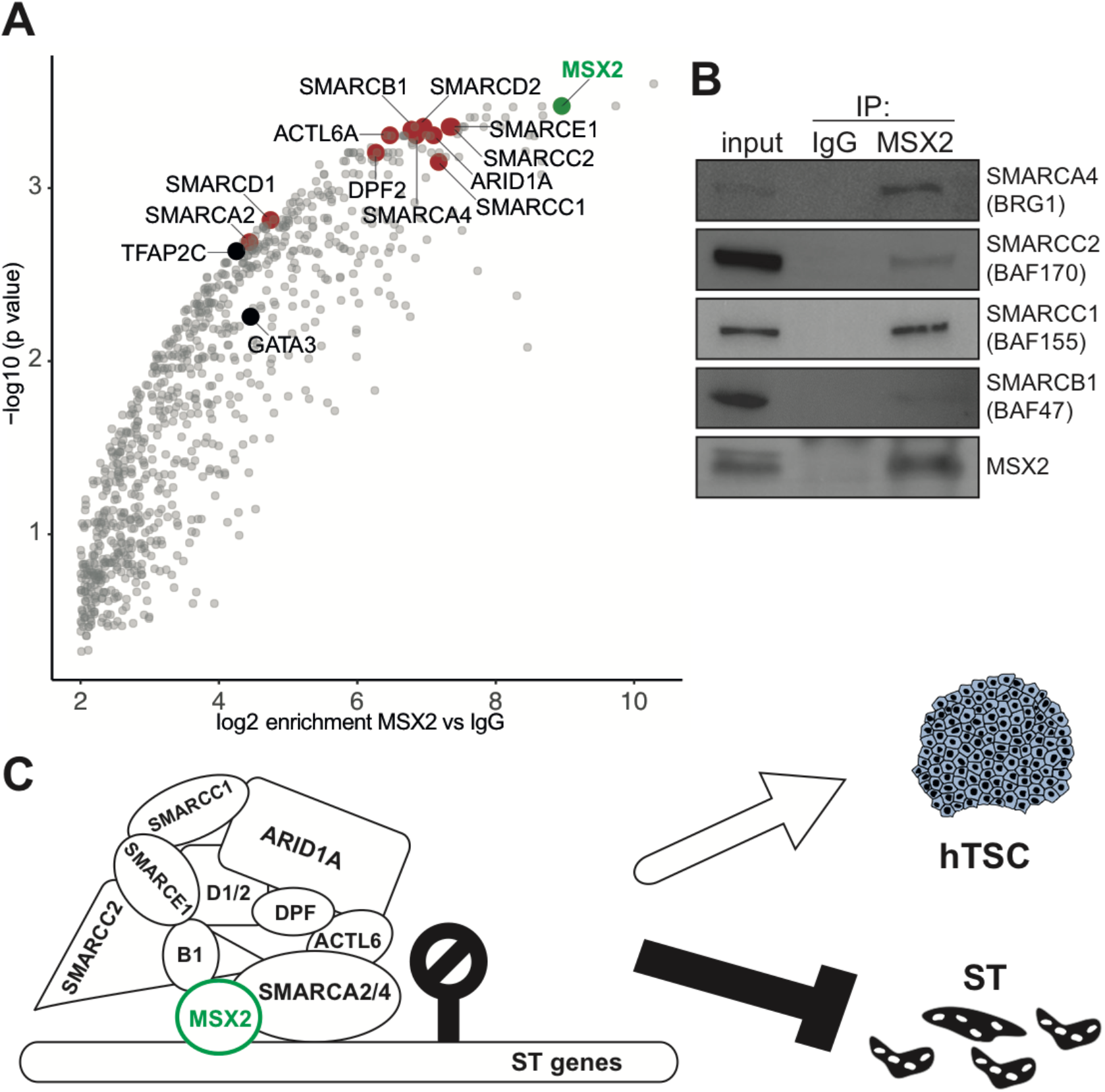
MSX2 cooperates with components of the SWI/SNF complex *(A)* MSX2 interactome identified in hTSC. Red and black mark selected MSX2 binding partners: components of SWI/SNF complex and prominent transcription factors, respectively. MSX2 is marked in green. The analysis is based on three biological replicates; IgG was used as control. (*B*) MSX2 immunoprecipitates analysed by Western blot probed with anti-SMARCA4 (BRG1), anti-SMARCC2 (BAF170), anti-SMARCC1 (BAF155) and anti-SMARCB1 (BAF47/INI1), confirming the prominent interaction between MSX2 and SWI/SNF complex components. *(C)* Model: MSX2 cooperates with the SWI/SNF complex and recruits it to the subset of ST-specific genes, to ensure their transcriptional silencing. Consequently, the ST state is blocked while the progenitor (hTSC) state is reinforced. ST: syncytiotrophoblast, hTSCs: human trophoblast stem cells.

In summary, we identified MSX2 as a central (core) regulator of trophoblast cell fate identity. We showed that while depletion of MSX2 activates ST transcriptional program, MSX2 forced expression blocks it. Importantly, a significant subset of these genes is directly bound and regulated by MSX2. Thus, we propose a mechanism, whereby MSX2 cooperates with the SWI/SNF complex and recruits it to the subset of ST-specific genes, to ensure transcriptional silencing and reinforcement of the progenitor state (Fig.4C). Our findings place MSX2 at the centre of the trophoblast specification process and uncover its vital role in the context of placental development and function.

## Materials and Methods

### Cell culture

hTSC lines CT30 and CT27 were a generous gift from Dr Hiroaki Okae (Tohoku University, Japan). The cells were cultured, passaged and differentiated as described (Okae et al. 2018) with minor modifications. For details see Supplemental Material.

### Genetic modifications

Lentiviral knock-down was carried out as described (Latos et al. 2015b), for details see Supplemental Material. To generate the inducible MSX2 expression construct, we cloned the coding sequence of MSX2 including the 3xFlag tag into PiggyBac-Tre-Dest-rTA-HSV-neo (a kind gift from Dr Joerg Betschinger, FMI Basel). After transfection, the hTSC were selected with 300 μg/ml G418 and expression was induced with 1 μg/ml doxycycline.

### Protein isolation and Western blot analyses

Whole cell lysates were prepared with TG buffer (20mM Tris-HCl pH 7.5, 137mM NaCl, 1mM EGTA, 1% Triton X-100, 10% glycerol and 1.5mM MgCl2). Following primary antibodies were used: anti-MSX2 (HPA005652, Sigma, 1:800), anti-human Chorionic Gonadotropin (CBG, A0231, Dako, 1:1000), anti-ENDOU (HAP067448, Sigma, 1:1000), anti-tubulin (ab6160, Abcam, 1:2000), anti-FlagM2 (F1804, Sigma, 1:2000), anti-GCM1 (HPA011343, Sigma, 1:1000).

### RT-qPCR

RNA was extracted using RNeasy mini kit (Qiagen) and treated with DNaseI (Qiagen) according to the manufacturer’s protocol. cDNA was synthesized using 1.5-3 μg RNA primed with random hexamers according to the RevertAid Reverse Transcriptase protocol (Thermo Scientific EP0442). DNA was diluted and qPCR performed using GoTaq qPCR Master Mix (A6002, Promega). Results are shown as means of indicated number of biological replicates (n) +/− S.E.M. Statistical significance was determined using a two-tailed, unpaired t-test. ****p<0.0001, ***p<0.001, **p<0.01, *p<0.05, ns: not significant. Primer sequences are provided in the Supplemental Material.

### RNA-seq

RNA was extracted using RNeasy mini kit (Qiagen) and treated with DNaseI (Qiagen). Indexed libraries were prepared with 500 ng RNA using QuantSEQ 3’ mRNA-Seq Library Prep Kit FWD for Illumina (015.96, Lexogen) according to manufacturer’s recommendations. The libraries were pooled and sequenced with a 100-base-pair single-end (MSX2_iOX) and 50-base-pair single-end protocol on Illumina HiSeq 2500 sequencer. For details of the bioinformatic analysis see Supplemental Material.

### Immunofluorescence

Placental tissue (7^th^ week of gestation) was embedded in paraffin and processed as described (Haider et al. 2016). Utilization of tissues and all experimental procedures were approved by the Medical University of Vienna ethics boards (Nr. 084/2009) and required written informed consent.

Cells were fixed in 4% paraformaldehyde/PBS for 20 min at 4°C, permeabilized and blocked for 30 min in 4% donkey serum and 0.1% Triton X-100 in PBS. The following primary antibodies with given dilutions were used: anti-MSX2 (HPA005652, Sigma, 1:250), anti-human Chorionic Gonadotropin (hCG, A0231, Dako, 1:300), anti-ENDOU (HAP067448, Sigma, 1:250). Alexa Fluor-conjugated secondary antibodies (A-21206, A-21207, Invitrogen) were applied at 1:1000 in 4% donkey serum and 0.1% Tween-20 in PBS blocking solution. Cells were counterstained with 4,6-diamidino-2-phenylindole (DAPI) and imaged using Zeiss Imager A2 microscope with Zen 2012 software.

### RIME

Rapid immunoprecipitation mass spectrometry of endogenous proteins (RIME) was carried out as described, using anti-MSX2 (HPA005652, Sigma) antibody (Mohammed et al. 2016).

### Co-immunoprecipitation

Cells were harvested and resuspended in Hunt Buffer (20mM Tris-HCl pH 8.0, 100mM NaCl, 1mM EDTA, 0.5% NP-40), followed by three freeze-thaw cycles for whole protein extraction. Protein G magnetic Dynabeads (10004D, Invitrogen) were pre-blocked with 1mg/ml BSA and 5 μg of rabbit anti-MSX2 (HPA005652, Sigma) or rabbit normal IgG (NI01, Sigma) were conjugated to the beads for 1h at RT. Affinity-purification by pre-bound beads was performed with 1000 μg of proteins overnight at 4°C with rotation. The next morning, beads were washed in Hunt Buffer and eluted. The following antibodies were used: anti-MSX2 (HPA005652, Sigma, 1:800), anti-BRG1 (sc-17796, Santa-Cruz, 1:1000), anti-BAF170 (sc-17838, Santa-Cruz, 1:500), anti-BAF155 (sc-48350, Santa-Cruz, 1:500), anti-BAF47 (sc-166165, Santa-Cruz, 1:250).

### ChIP-seq

Immunoprecipitations were carried out as described (Latos et al. 2015a) using anti-MSX2 (HPA005652, Sigma) antibody, for details, including the bioinformatic analysis see the Supplemental Material.

## Competing Interest Statement

The authors declare no conflict of interest

## Acknowledgments

This work was supported by the Austrian Science Fund (grant P-31738-B26 awarded to PL and MK). Sequencing was performed at the Vienna Biocenter Core Facilities Next Generation Sequencing Unit (www.viennabiocenter.org/facilities). We are grateful to Sasha Mendjan and Malte Mederacke for valuable comments.

## Author contributions

RH and PL conceived the study, performed experiments and analysed the data. AL contributed to the bioinformatic analysis. KM performed mass spectrometry analysis. SH and MK provided and processed placental tissue. PL wrote the manuscript.

## Notes

### Competing Interest Statement

The authors have declared no competing interest.

