## Supplemental Material for "MSX2 safeguards syncytiotrophoblast fate of human trophoblast stem cells"

Hornbachner et al.,

List of Material provided:

- Supplemental Fig.S1: Additional evidence that depletion of MSX2 results in loss of silencing of syncytiotrophoblast genes.
- Supplemental Fig.S2: Additional evidence that ectopic expression of MSX2 blocks syncytiotrophoblast cell fate.
- Supplemental Fig.S3: Additional data on MSX2 binding and silencing of syncytiotrophoblast genes in hTSC.
- Supplemental Fig.S4: Additional data on MSX2-SWI/SNF complex cooperation.
- Additional details on Materials & Methods

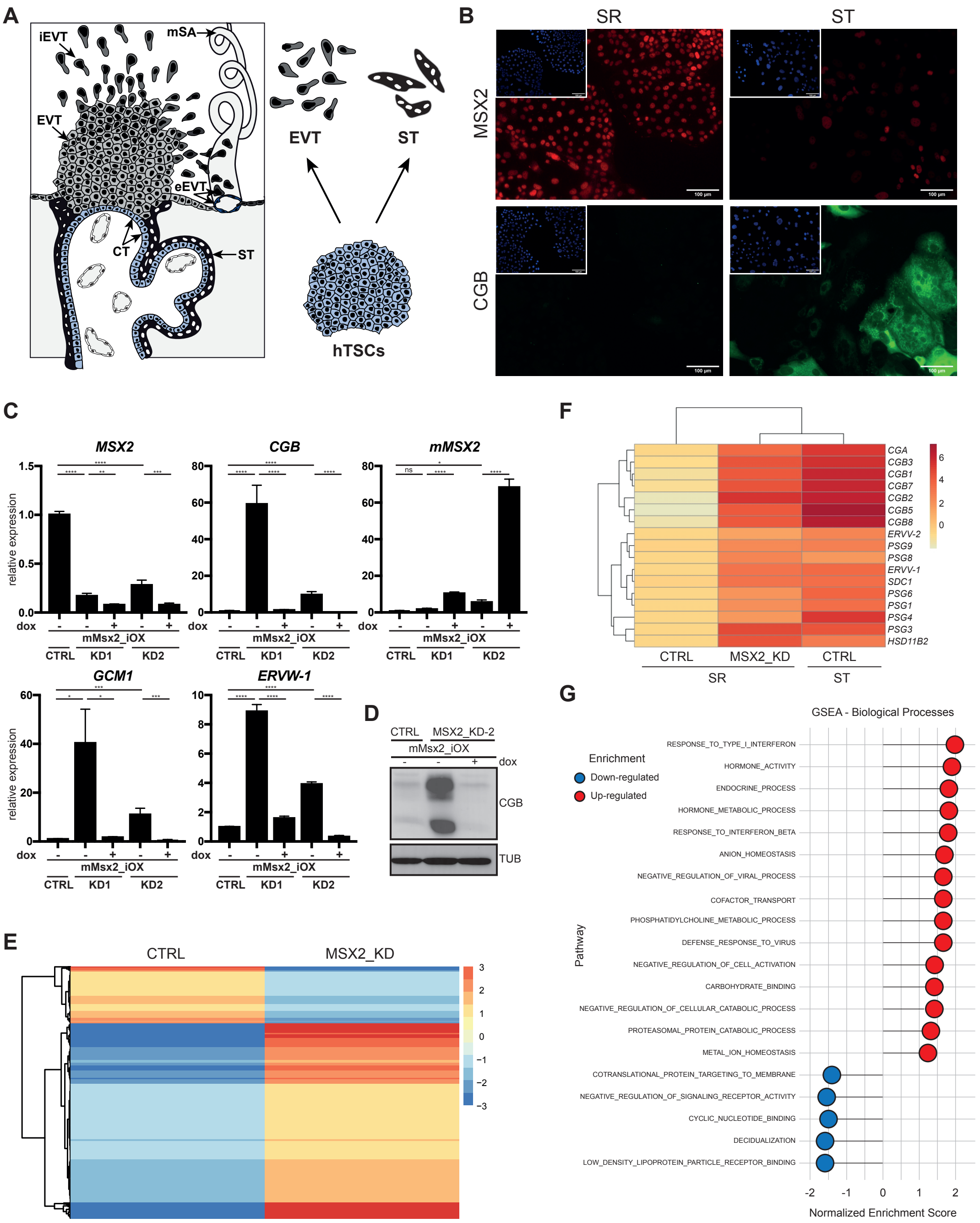
A
iEVT
mSA
EVT
eEVT
CT
ST
EVT
ST
hTSCs
B
SR
ST
MSX2
CGB
C
MSX2
CGB
mMSX2
relative expression
dox
mMsx2_iOX
CTRL
KD1
KD2
GCM1
ERVW-1
D
CTRL
MSX2_KD-2
mMsx2_iOX
dox
CGB
TUB
E
CTRL
MSX2_KD
F
CGA
CGB3
CGB1
CGB7
CGB2
CGB5
CGB8
ERVV-2
PSG9
PSG8
ERVV-1
SDC1
PSG6
PSG1
PSG4
PSG3
HSD11B2
CTRL
MSX2_KD
CTRL
SR
ST
G
GSEA - Biological Processes
Enrichment
Down-regulated
Up-regulated
Pathway
RESPONSE_TO_TYPE_I_INTERFERON
HORMONE_ACTIVITY
ENDOCRINE_PROCESS
HORMONE_METABOLIC_PROCESS
RESPONSE_TO_INTERFERON_BETA
ANION_HOMEOSTASIS
NEGATIVE_REGULATION_OF_VIRAL_PROCESS
COFACTOR_TRANSPORT
PHOSPHATIDYLCHOLINE_METABOLIC_PROCESS
DEFENSE_RESPONSE_TO_VIRUS
NEGATIVE_REGULATION_OF_CELL_ACTIVATION
CARBOHYDRATE_BINDING
NEGATIVE_REGULATION_OF_CELLULAR_CATABOLIC_PROCESS
PROTEASOMAL_PROTEIN_CATABOLIC_PROCESS
METAL_ION_HOMEOSTASIS
COTRANSLATIONAL_PROTEIN_TARGETING_TO_MEMBRANE
NEGATIVE_REGULATION_OF_SIGNALING_RECEPTOR_ACTIVITY
CYCLIC_NUCLEOTIDE_BINDING
DECIDUALIZATION
LOW_DENSITY_LIPOPROTEIN_PARTICLE_RECEPTOR_BINDING
Normalized Enrichment Score

**Supplemental Figure S1**

Depletion of MSX2 results in loss of silencing of syncytiotrophoblast genes.

*(A)* Schematic representation of a fragment of a villous tree in human placenta with main trophoblast cell types marked and their *in vitro* counterpart derivatives. EVT: extravillous trophoblast, iEVT: interstitial EVT, eEVT: endovascular EVT, ST: syncytiotrophoblast, CT: cytotrophoblast, mSA: maternal spiral artery, hTSCs: human trophoblast stem cells.

*(B)* Immunostaining of self-renewing (SR) and syncytiotrophoblast differentiated (ST) hTSCs for MSX2 and CGB. ST: syncytiotrophoblast, CT: cytotrophoblast.

*(C)* RTQPCR analysis of control (CTRL) and MSX2-depleted (MSX2_KD1 and MSX2_KD2) hTSCs carrying a doxycycline (dox) inducible mouse Msx2 transgene (mMsx2_iOX) and cultured in self-renewal conditions in the presence (+dox) or absence (-dox) of dox. The bars represent a mean of three biological replicates (n=3) with S.E.M., expression in CTRL was set to 1. ****$p<0.0001$, ***$p<0.001$, **$p<0.01$, *$p<0.05$, ns: not significant. MSX2 indicates expression of the human, endogenous MSX2, while mMsx2 of the mouse Msx2 transgene.

*(D)* Western blot analysis for CGB of control (CTRL) and MSX2-depleted (MSX2_KD2) hTSCs carrying a doxycycline (dox) inducible mouse Msx2 transgene (mMsx2_iOX), cultured in self-renewal conditions in the presence (+dox) or absence (-dox) of dox. Tubulin (TUB) serves as a loading control.

*(E)* Hierarchical clustering of genome-wide RNA profiles of MSX2-depleted (MSX2_KD: MSX2_KD1 (n=3) and MSX2_KD2 (n=1)) and control (CTRL (n=3) hTSCs cultured in self-renewal conditions. Analysis ($|log2FC|>1$, adj $p<0.05$) revealed 522 up-regulated and 152 down-regulated genes in MSX2_KD lines compared to control.

*(F)* Clustered heat map depicting expression of selected syncytiotrophoblast markers in control (CTRL) and MSX2-depleted (MSX2_KD) hTSC lines cultured in self-renewal conditions (SR) (details as in A.) and, as comparison, in cells differentiated for 6 days to syncytiotrophoblast (ST).

*(G)* Gene Set Enrichment Analysis (GSEA) for biological processes of de-regulated genes in MSX2_KD compared to control (CTRL), based on RNA-Seq analysis as in Supplemental Fig.1E.

Hornbachner et al._Supplemental_Fig_S2

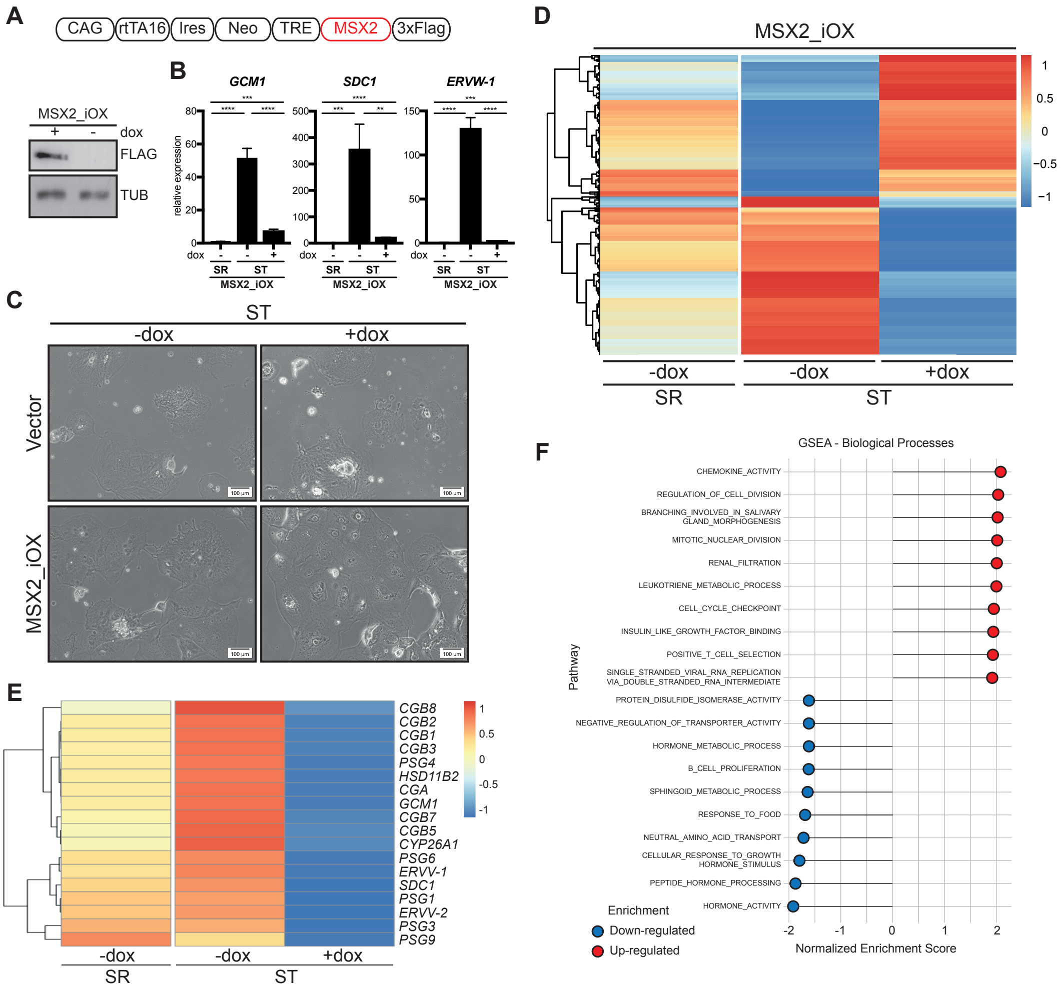

**Supplemental Figure S2**

Ectopic expression of MSX2 blocks syncytiotrophoblast cell fate.

*(A)* Schematic depiction of a doxycycline (dox) inducible construct carrying a 3xFlag-tagged coding sequence of human MSX2 and a Western blot showing expression of this transgene upon 48h of dox treatment.

*(B)* RTQPCR analysis for syncytiotrophoblast markers *GCM1*, *SDC1* and *ERVW-1* in hTSC line carrying a doxycycline (dox) inducible MSX2 transgene and differentiated to syncytiotrophoblast (ST) in the presence (+dox) or absence (-dox) of dox. The bars represent an average of three independent biological replicates (n=3) with S.E.M.; expression in self-renewal conditions (SR) was set to 1. ****$p<0.0001$, ***$p<0.001$, **$p<0.01$, *$p<0.05$, ns: not significant.

*(C)* Bright field images of hTSC lines carrying a doxycycline (dox) inducible MSX2 transgene (MSX2_iOX) or an empty vector (Vector) and differentiated for 6 days into syncytiotrophoblast (ST) in the presence (+dox) or absence (-dox).

*(D)* Hierarchical clustering of genome-wide expression profiles of hTSC carrying a doxycycline (dox) inducible MSX2 transgene (MSX2_iOX) differentiated for 6 days into syncytiotrophoblast (ST) in the presence (+dox) or absence (-dox) of doxycycline. Analysis ($|\log2FC|>1$, adj $p<0.05$) revealed 1285 up-regulated and 1433 down-regulated genes in MSX2_iOX (+dox) compared to (-dox) control. For comparison, levels of these differentially expressed genes in -dox, self-renewal conditions (SR) are displayed.

*(E)* Clustered heat map depicting expression of selected syncytiotrophoblast markers as in Supplemental Fig. S2D.

*(F)* Gene Set Enrichment Analysis (GSEA) for biological processes of de-regulated genes in MSX2_iOX ST (+dox) in comparison to MSX2_iOX ST (-dox), based on RNA-Seq analysis as in Supplemental Fig. S2D.

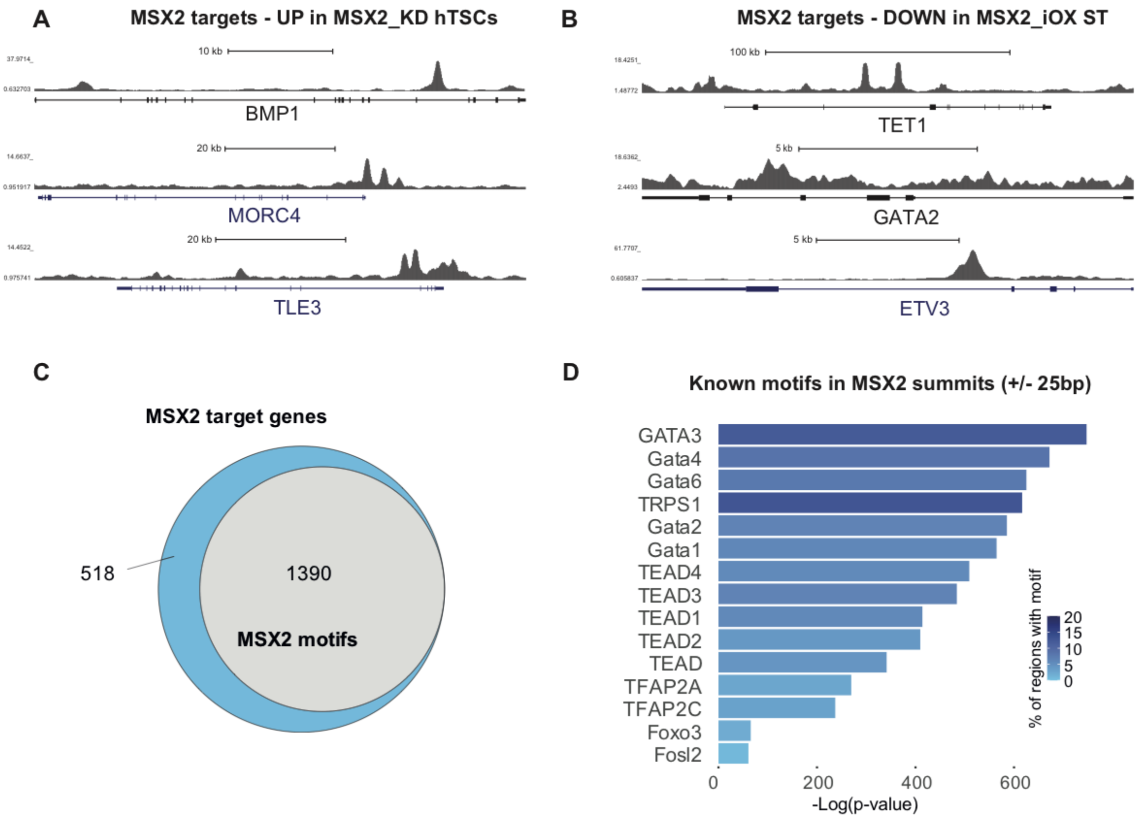


**Supplemental Figure S3** MSX2 binds and silences the syncytiotrophoblast genes in hTSCs.

*(A)* UCSC genome browser snapshots for MSX2 targets identified among the upregulated genes in hTSCs upon MSX2_KD.

*(B)* UCSC genome browser snapshots for MSX2 targets identified among the downregulated genes in ST upon MSX2 overexpression (MSX2_iOX ST).

*(C)* Number of MSX2 target genes with the MSX2 motif, identified by HOMER *de novo* motif detection. Total targets, blue; MSX2 motif containing targets, grey.

*(D)* Bar graph showing the identified known TF motifs around the MSX2 peak summits. The bar length represents -Log(*p*-value) of enrichment and the colour scale indicates percentage of regions containing the respective TF motif.

**A**

| | | | | MSX2 | | | | | | IgG | | | | | |
|---|---|---|---|---|---|---|---|---|---|---|---|---|---|---|---|
| **Gene** | **Alias** | **log2FC** | **p-value** | **# unique peptides** | | | **% coverage** | | | **# unique peptides** | | | **% coverage** | | |
| **MSX2** | MSX2 | 8.95 | 6.33E-07 | 3 | 3 | 2 | 22.5 | 22.1 | 16.9 | 0 | 0 | 0 | 0.0 | 0.0 | 0.0 |
| **SMARCE1** | BAF57 | 7.36 | 3.28E-06 | 10 | 10 | 11 | 28.7 | 26.5 | 32.9 | 0 | 0 | 0 | 0.0 | 0.0 | 0.0 |
| **SMARCC2** | BAF170 | 7.33 | 3.97E-06 | 21 | 23 | 20 | 25.6 | 28.6 | 24.1 | 3 | 4 | 0 | 2.6 | 3.5 | 0.0 |
| **SMARCC1** | BAF155 | 7.18 | 2.59E-05 | 23 | 31 | 31 | 34.3 | 37.5 | 35.5 | 0 | 2 | 0 | 1.7 | 4.9 | 0.0 |
| **ARID1A** | BAF250A | 7.10 | 7.10E-06 | 31 | 31 | 42 | 20.1 | 20.0 | 24.6 | 0 | 1 | 0 | 0.0 | 0.0 | 0.0 |
| **SMARCD2** | BAF60B | 6.95 | 4.45E-06 | 18 | 13 | 15 | 43.9 | 34.3 | 38.4 | 0 | 0 | 0 | 0.0 | 0.0 | 0.0 |
| **SMARCA4** | BRG1 | 6.85 | 7.48E-06 | 12 | 17 | 20 | 16.8 | 20.3 | 19.7 | 0 | 0 | 0 | 0.0 | 0.0 | 0.0 |
| **SMARCB1** | BAF47 | 6.78 | 5.76E-06 | 11 | 12 | 7 | 33.5 | 42.1 | 27.5 | 0 | 0 | 0 | 0.0 | 0.0 | 0.0 |
| **ACTL6A** | BAF53A | 6.47 | 8.01E-06 | 11 | 11 | 9 | 32.4 | 37.5 | 25.4 | 0 | 0 | 0 | 0.0 | 0.0 | 0.0 |
| **DPF2** | BAF45D | 6.27 | 1.79E-05 | 8 | 8 | 11 | 23.0 | 31.0 | 32.5 | 0 | 0 | 0 | 0.0 | 0.0 | 0.0 |
| **SMARCA2** | BRM | 4.44 | 1.66E-04 | 9 | 10 | 9 | 16.0 | 15.6 | 13.3 | 0 | 0 | 0 | 0.0 | 0.0 | 0.0 |
| **SMARCD1** | BAF60A | 4.74 | 9.02E-05 | 9 | 10 | 9 | 23.9 | 28.2 | 24.3 | 0 | 0 | 0 | 0.0 | 0.0 | 0.0 |

**Supplemental Figure S4** MSX2 cooperates with components of the SWI/SNF complex.

*(A)* Details of mass spectrometry identification of the SWI/SNF complex components from Fig.4A.

**Additional details on Materials & Methods**

*Chromatin immunoprecipitation*

Cells (1-2x$10^8$) were fixed in 2mM Di(N-succinimidyl) glutarate (DSG) (80424, Sigma) in PBS at room temperature (RT) for 45 min. After washing with PBS, cells were fixed again in 1% formaldehyde (28908, Sigma) in TS base media at RT for 12 min. Fixation was stopped by adding glycine to a final concentration of 0.125 M. Cells were washed twice with PBS and resuspended in wash buffer 1 (10mM Hepes pH7.5, 10mM EDTA, 0.5mM EGTA and 0.75% Triton X-100) and incubated at 4C for 10min. After pelleting, cells were resuspended in wash buffer 2 (10mM Hepes pH7.5, 200mM NaCl, 1mM EDTA and 0.5mM EGTA) and incubated at 4°C for 10min. After pelleting, cells were lysed in the lysis/sonication buffer (150mM NaCl, 25mM Tris pH 7.5, 5mM EDTA, 0.1%Triton, 1% SDS and 0.5% sodium deoxycholate) with cOmplete™ EDTA-free Protease Inhibitor Cocktail (11873580001, Sigma) on ice for 30min. Chromatin was sonicated 15s on /30s off for 23-25 cycles using the UW2070 sonicator (Bandelin) to the average 300-bp fragments. Chromatin was diluted 1/10 with the dilution buffer (150mM NaCl, 25mM Tris pH 7.5, 5mM EDTA, 1% Triton X-100, 0.1% SDS and 0.5 % sodium deoxycholate) containing complete protease inhibitors. Protein G magnetic Dynabeads (10004D, Invitrogen) were blocked with 1mg/ml BSA and tRNA at 4°C for 1h and washed with buffer A (150mM NaCl, 25mM Tris pH 8.0, 1mM EDTA, 1% NP-40, 0.1 % SDS and 0.5 % sodium deoxycholate). Chromatin was pre-cleared with pre-blocked beads at 4°C for 1h. Four hundred micrograms of chromatin and seven micrograms of antibody rabbit anti-MSX2 (HPA005652, Sigma) and rabbit normal IgG (NI01, Sigma) were used per IP. IP was performed overnight at 4°C with rotation. Pre-blocked magnetic beads were added next morning for 7-8 h. Beads were washed at 4°C with buffer A three times, buffer B (50mM Tris pH 8.0, 500mM NaCl, 0.1% SDS, 0.5 % sodium deoxycholate, 1% NP-40 and 1mM EDTA), buffer C (50mM Tris pH 8.0, 250mM LiCl, 0.5% sodium deoxycholate, 1% NP-40 and 1mM EDTA) and rinsed with TE buffer. DNA was eluted from beads in the elution buffer (1% SDS, 0.1M $NaHCO_3$). Samples were treated with RNAse A and Proteinase K and reverse-crosslinked overnight by incubation for 2h at 37C, followed by 12h at 65C. DNA was purified on the PCR purification columns (Qiagen). To generate a library, DNA from 2-4 IPs was pooled and the NEBNext Ultra II DNA library preparation master mix (New England Biolabs, E7645) was used according to the manufacturer's instructions. Libraries were sequenced on an Illumina HiSeq 2500 sequencer with 50-base pair (bp) single-end protocol.

*Bioinformatic analysis*

RNA quantification, QuantSeq

Libraries generated with the QuantSeq FWD kit (Lexogen) were sequenced on an Illumina HiSeq2500v4 at single read 50bp. Reads were trimmed with Trim Galore! (0.6.5). Quality control was performed using fastQC (0.11.9) (Andrews). Transcripts were mapped to the hg38 human reference genome using STAR (2.7.3a) (Dobin et al. 2013). After indexing with samtools (1.10) reads in genes were counted using Rsubread (2.0.0) (Liao et al. 2019). Differential expression analysis was conducted using DESeq2 (1.24.0) (Love et al. 2014). MA plot was generated with ggplot2 (3.3.3) and the heatmap using pheatmap (1.0.12) (Kolde). Gene set enrichment analysis (GSEA) was performed using fgsea (1.10.1) (Korotkevich et al.) with biological process GO annotations downloaded from Molecular Signatures Database (MSigDB).

*Chromatin Immunoprecipitation Sequencing*

Libraries were sequenced on the Illumina HiSeq2500 v4 at single read 50bp aiming for a sequencing depth of 40 million reads/sample. Raw reads were trimmed with CUTADAPT (2.8) and aligned to the human reference genome GRCh38 with bowtie 2 (2.3.5.1) (Langmead and Salzberg 2012). Alignments were further processed using samtools (1.10). Peaks were called using MACS2 (2.2.7.1) with default parameters taking uniquely mapped reads with a merged input as the control (Zhang et al. 2008). 2661 high confidence peaks were generated with estimate_idr1d of the R package idr2d (1.4.0), using a cut-off of FDR<0.05 across all samples (Krismer et al. 2020). Feature distribution was determined, and peaks were annotated with ChIPseeker (1.26.0) with following priorities: Promoter, 5’UTR, 3’UTR, Exon, Intron, Downstream, Intergenic (Yu et al. 2015). For visualization we generated RPKM normalized coverage files with deeptools (3.5.0) and bamCoverage (Ramírez et al. 2016). *De novo* motif search and motif enrichment analyses were conducted with HOMER using the summits of all replicates included in the significantly reproducible peaks. MSX2 motif search was done on the IDR-validated peak set by HOMER. Gene annotations of the MSX2 peaks containing an MSX2 motif were derived from ChIPseeker (see above).

*Genetic modifications*

To deplete MSX2, we generated pLKO.1-based constructs containing either shRNA against MSX2 (TRCN0000234848 and TRCN0000234849) or a shRNA against GFP as a control by cloning annealed oligos into the AgeI/EcoRI sites of pLKO.1-neo (13425, Addgene). The plasmids were simultaneously transfected into HEK293T cells: pLKO.1 (individual constructs containing different shRNA), psPAX2 (encodes Gag and Pol sequences to package the lentivirus), and pMD2.G (encodes the G protein envelope protein) using lipofectamine 3000 (Thermo Fisher Scientific). Supernatant was collected after 48h and used to transduce hTSC for a minimum of 16h. Cells were selected using 300 µg/ml G418 (A1720, Sigma).

*Cell culture conditions*

Briefly, the cells were cultured in the basal media (DMEM/F12 (11320074, Thermo Fisher), 1x ITS-X (51500056, Thermo Fisher), 0.2% FBS (PAA), 1.5 µg/mL L-ascorbic acid (A4403, Sigma)) supplemented with 2 µM CHIR99021 (2520691, Peprotech), 50ng/mL EGF (AF-100-15, Peprotech), 0.5 µM A83-01 (9094360, Peprotech), and 5µM Y27632 (1293823, Peprotech), on dishes coated with 10µg/ml Fibronectin (FC0101, Sigma). Cells were passaged every 3-4 days using TrypLE (12605036, Thermo Scientific). ST differentiation was induced in basal medium supplemented with 2 µM Forskolin (6652995, Peprotech), 5 µM Y27632 and 4% KnockOut Serum Replacement (10828010, Thermo Scientific).

*Mass Spectrometry*

Liquid Chromatography mass spectrometry (LC-MS) was performed on an UltiMate 3000 RSLC nano system (Thermo Scientific) coupled to a Q Exactive HF-X mass spectrometer (Thermo Scientific) and equipped with a Nanospray Flex Ion Source (Thermo Scientific). For peptide identification, the RAW-files were processed using Proteome Discoverer (2.3.0.523) (Thermo Scientific). The MS/MS spectra were searched against the Uniprot human proteome database 2020-10-12 (20,536 sequences; 11,395,384 residues) and the default list of common contaminants using MSAmanda (v2.0.0.12368) (Dorfer et al. 2014). Beta-methylthiolation on cysteine was set as a fixed modification. Localization of post-translational modification sites within the peptides was performed using the function ptmRS of phospho-RS (Taus et al. 2011). The maximal number of missed cleavages was set to 2, using tryptic enzymatic specificity. The results were filtered to 1 % FDR on the protein level using the Percolator algorithm integrated in Proteome Discoverer. Peptide areas were quantified using apQuant (Doblmann et al. 2019).

*Primer sequences*

| **Name** | **Sequence** |
|---|---|
| RTQPCR | |
| hPBGD-1F | GGAGCCATGTCTGGTAACGG |
| hPBGD-1R | CCACGCGAATCACTCTCATCT |

| | |
|---|---|
| hMSX2-1F | GCAGGAACCCGGCCGATATT |
| hMSX2-1R | CTGACGGAACTTGCGCTCCA |
| hCGB-F | CAGCATCCTATCACCTCCTGGT |
| hCGB-R | CTGGAACATCTCCATCCTTGGT |
| hGCM1-1F | GCTGGGACTTGAACCAGCAGT |
| hGCM1-1R | CTGGATCGGCCCACTCAAGC |
| hSDC1_F | CTATTCCCACGTCTCCAGAACC |
| hSDC1_R | GGACTACAGCCTCTCCCTCCTT |
| ERVW-1_1F | CTACCCCAACTGCGGTTAAA |
| ERVW-1_1R | GGTTCCTTTGGCAGTATCCA |
| hTEAD4_F | GCCAGTCCAGCCCAAGCTAC |
| hTEAD4_R | CATTGGAGGGTCCCCGTTCG |
| shRNAs | |
| CTRL | GCAAGCTGACCCTGAAGTTCAT |
| MSX2_KD1 | AGCGCAAGTTCCGTCAGAAAC |
| MSX2_KD2 | TGCAGGCAGCGTCCATATATG |

*Supplemental References*

Andrews S. FastQC: A quality control tool for high throughput sequence data. https://www.bioinformatics.babraham.ac.uk/projects/fastqc/.

Dobin A, Davis CA, Schlesinger F, Drenkow J, Zaleski C, Jha S, Batut P, Chaisson M, Gingeras TR. 2013. STAR: ultrafast universal RNA-seq aligner. *Bioinformatics* **29**: 15–21.

Doblmann J, Dusberger F, Imre R, Hudecz O, Stanek F, Mechtler K, Dürnberger G. 2019. apQuant: Accurate Label-Free Quantification by Quality Filtering. *J Proteome Res* **18**: 535–541.

Dorfer V, Pichler P, Stranzl T, Stadlmann J, Taus T, Winkler S, Mechtler K. 2014. MS Amanda, a universal identification algorithm optimized for high accuracy tandem mass spectra. *J Proteome Res* **13**: 3679–3684.

Kolde R. pheatmap: Pretty Heatmaps. https://CRAN.R-project.org/package=pheatmap.

Korotkevich G, Sukhov V, Sergushichev. Fast gene set enrichment analysis.

Krismer K, Guo Y, Gifford DK. 2020. IDR2D identifies reproducible genomic interactions. *Nucleic Acids Res* **48**: e31–e31.

Langmead B, Salzberg SL. 2012. Fast gapped-read alignment with Bowtie 2. *Nat Methods* **9**: 357–359.

Liao Y, Smyth GK, Shi W. 2019. The R package Rsubread is easier, faster, cheaper and better for alignment and quantification of RNA sequencing reads. *Nucleic Acids Res* **47**: e47–e47.

Love MI, Huber W, Anders S. 2014. Moderated estimation of fold change and dispersion for RNA-seq data with DESeq2. *Genome Biol* **15**: 550–21.

Ramírez F, Ryan DP, Grüning B, Bhardwaj V, Kilpert F, Richter AS, Heyne S, Dündar F, Manke T. 2016. deepTools2: a next generation web server for deep-sequencing data analysis. *Nucleic Acids Res* **44**: W160–5.

Taus T, Köcher T, Pichler P, Paschke C, Schmidt A, Henrich C, Mechtler K. 2011. Universal and confident phosphorylation site localization using phosphoRS. *J Proteome Res* **10**: 5354–5362.

Yu G, Wang L-G, He Q-Y. 2015. ChIPseeker: an R/Bioconductor package for ChIP peak annotation, comparison and visualization. *Bioinformatics* **31**: 2382–2383.

Zhang Y, Liu T, Meyer CA, Eeckhoute J, Johnson DS, Bernstein BE, Nusbaum C, Myers RM, Brown M, Li W, et al. 2008. Model-based analysis of ChIP-Seq (MACS). *Genome Biol* **9**: R137–9.
